# Field evidence for the role of plant volatiles induced by caterpillar-derived elicitors in the prey location behavior of predatory social wasps

**DOI:** 10.1101/2024.03.29.587343

**Authors:** Patrick Grof-Tisza, Ted C.J. Turlings, Carlos Bustos-Segura, Betty Benrey

## Abstract

1. One assumed function of herbivore-induced plant volatiles (HIPVs) is to attract natural enemies of the inducing herbivores. Field evidence for this is scarce and often indirect. Also, the assumption that elicitors in insect oral secretions that trigger the volatile emissions are essential for attraction of natural enemies has not yet been demonstrated under field conditions.
2. After observing social wasps removing caterpillars from maize plants in an agricultural field, we hypothesized that these wasps use HIPVs to locate their prey. To test this, we conducted an experiment that simultaneously explored the importance of caterpillar oral secretions in the interaction.
3. We found that *Spodoptera* caterpillars placed on mechanically damaged plants treated with oral secretion were more likely to be attacked by wasps compared to caterpillars on plants that were only mechanically wounded. Both of the the latter treatments were considerably more attractive than plants that were only treated with oral secretion or left untreated. Subsequent analyses of headspace volatiles confirmed differences in emitted volatiles that likely account for the differential predation events across the treatments.
4. These findings highlight the importance of HIPVs in prey location by social wasps and provide evidence for the role that elicitors play in inducing attractive odor blends.

## INTRODUCTION

Herbivore infestations can trigger substantial changes in the volatile compounds emitted by plants under attack (Hare, 2011). It is widely assumed that these herbivore-induced plant volatiles (HIPVs) serve as an indirect defense mechanism by attracting natural enemies of the attacking herbivore (reviewed in Dicke & Baldwin, 2010; Heil 2014; Pearse et al., 2020; Turlings & Erb, 2018). While numerous studies have shown that HIPVs can indeed attract predators and parasitoids (e.g., Dicke & Sabelis 1988; Dicke & van Loon, 2000; Kessler & Baldwin, 2001; Turlings et al., 1990), most have been conducted under controlled laboratory settings using olfactometers, wind tunnels, and observation arenas; relatively few studies have addressed this question under natural field conditions, as commonly noted in reviews (Dicke & Baldwin, 2010; Kessler & Baldwin, 2003; Takabayashi & Shiojiri, 2019). Those that have, largely focused on attraction of carnivorous arthropods and parasitoids without quantifying changes in herbivore abundance or parasitism rates (e.g., Simpson et al., 2011; Yu et al., 2008). Rarer still is evidence linking reductions of herbivores or increased parasitism to consequential changes in herbivory, (but see Cuny et al., 2018; de Lange et al., 2018; Fritzsche-Hoballah & Turlings, 2001; Gomez & Zamora, 1994; Kessler & Baldwin, 2001; Schuman et al., 2012). Consequently, despite the abundant and compelling laboratory studies demonstrating the attraction of natural enemies to HIPVs, our understanding of their true ecological importance and potential applications in pest management is limited.

Plants exhibit distinct responses to damage caused by biological agents such as herbivores or pathogens compared to mechanical damage (Bonaventure et al., 2011; Kessler & Baldwin, 2003). More notably, they can discern between herbivores employing different feeding modalities (Leitner et al., 2005; Turlings et al., 1998) and even among species of lepidopteran larvae (Ling et al., 2021), enabling precise and tailored responses to specific threats. This recognition can be achieved through herbivore elicitors (also known as herbivore-associated molecular patterns (Acevedo et al., 2015; Mithöfer & Boland, 2008). Elicitors are molecules from an herbivore present in feces, saliva, and oviposition secretions (Hilker & Meiners, 2010; Schmelz, 2015; Uemura & Arimura, 2019). Once recognized by the plants, elicitors trigger defense pathways such as those leading to the release or amplification of specific volatile compounds. The resulting induced volatile blend serves as a dependable cue for predators, aiding them in distinguishing between plants under attack by their preferred prey (Kessler & Baldwin, 2001). This distinction is particularly vital in the case of parasitoids, where the specific induced odor blend can signify the presence of suitable hosts (De Moraes et al., 1998).

Various elicitors (e.g., caeliferins, Alborn et al., 2007; β-glucosidase, Mattiacci et al., 1995; vitellogenin, Zeng et al., 2023), have been identified within the oral secretions of herbivores, including *Spodoptera* caterpillars (volicitin, Alborn et al., 1997; inceptin, Schmelz et al., 2006). Simulated herbivory experiments have consistently revealed rapid emission of terpenoids and other important volatiles involved plant-insect signaling following the application of *Spodoptera* caterpillar oral secretions to mechanically inflicted wounds compared to plants that were only subjected to mechanical wounding (Turlings et al., 1993, 2000). These findings suggest that factorial experiments should be conducted to disentangle the relative importance of factors contributing to the cues, including HIPVs, that play a role in the recruitment of natural enemies (Mamin et al., 2023; Waterman et al., 2019). While this experimental methodology is commonplace in laboratory experiments (e.g., Arce et al., 2021; Bricchi et al., 2010; Schittko et al., 2001), field studies have often relied on natural herbivory, potentially confounding the identification of key factors driving predator and parasitoid attraction (but see Bernasconi Ockroy et al., 2001; Schuman et al. 2012). For a better understanding of the ecological implications of volatile-mediated interactions between herbivores and their natural enemies and to leverage top-down pest regulation in support of sustainable agriculture, targeted manipulative studies under realistic natural and agroecological conditions are warranted.

The motivation for this current study came through observations of social predatory wasps removing *Spodoptera frugiperda* caterpillars from maize plants within an agricultural field in Oaxaca, Mexico (Grof-Tisza et al. 2023). We hypothesized that these wasps used HIPVs to locate their prey. While it is known that predatory wasps use both visual and olfactory cues to locate their prey (Richter, 2000), few studies have investigated the role of volatiles from prey or prey-infested plants in their foraging behavior (Aldrich et al., 1985; Cornelius, 1993; M; Saraiva et al., 2017) with one laboratory study analyzing the attraction of HIPVs using a y-tube olfactometer (Saraiva et al., 2017). It is unknown to what extent social wasps rely on HIPVs during foraging under natural conditions, which is also of applied importance in light of a growing interest in the use of these predators in biocontrol (Montefusco et al., 2017; Prezoto et al., 2019a; Southon et al., 2019).

To investigate the importance of olfaction and HIPVs in the foraging behavior of predatory social wasps, we performed a sentinel prey experiment in a maize field using potted maize plants and two armyworm caterpillar species, the fall armyworm (*Spodoptera frugiperda,* hereafter FAW) and the velvet armyworm (*Spodoptera latifasica,* hereafter VAW). Using a factorial design, we manipulated factors that induce plant volatiles: the mechanical wounding of leaves and caterpillar oral secretions that contain elicitors. We then examined their influence on the attack rate of FAW and VAW caterpillars by naturally occurring wasps under field conditions. Secondly, we analyzed emitted volatiles from maize plants subjected to the same treatments under laboratory conditions. The latter aimed to provide evidence that observed differences in attack rates between the treatments in the sentinel field experiments were driven by differences in inducible odors.

## METHODS

### Sentinel Prey Experiment

#### Field Site

The study was conducted over two seasons (2022 and 2023) in a 925 m^2^ agricultural field near Bajos de Chila Oaxaca, Mexico (15,91818°N, 97,16401°W) within an experimental plantation of Milpa (Grof-Tisza et al. 2023), a traditional Mesoamerican intercropping system comprised of maize, squash, and beans, (*Zea mays*, Mexican hybrid; var White maize, *Cucurbita argyrosperma* ssp. argyrosperma; *Phaseolus vulgaris*). The field was surrounded by mango trees (*Magnifera* sp.) and coconut palms (*Cocos* sp.), which provided natural habitats for wasp nests (e.g., *Polistes* sp., *Polybia* sp.). Social wasps are voracious predators consuming upwards of 4,000 prey items, primarily lepidopteran larvae, during their brief adult life span (Prezoto et al., 2006; Prezoto et al., 2007).

#### Plants and Insects

Three to four maize seeds of the same local variety that were planted in the experimental field were sown in 1.5 L plastic pots containing field collected soil. To prevent herbivory, plants were grown in large mesh cages (2 x 2 x 4 m; BioQuip, CA, USA) for 10-12 days at our field site prior to the experiment. The two healthiest plants of equivalent size in each pot were left for the experiments; remaining plants were removed before imposing treatments two weeks after germination.

Caterpillars of FAW and VAW were collected from maize and bean plants at the same field site and reared in our field laboratory under natural light and humidity conditions and fed an artificial diet (Beet Armyworm diet, Frontier Agri Science USA). These two species are commonly found in the study area, feed on maize, and are readily attacked by predatory wasps (Grof-Tisza et. 2023). Only FAW caterpillars (third instar) were used as a source of oral secretion, which is known to elicit HIPV emissions in maize (reviewed in Turlings and Erb, 2018), the caterpillars were switched to a maize diet 8-16 hours prior to using them in the field experiment. Third instar caterpillars of both species were used as sentinel prey.

#### Field Trials

Pots containing two maize plants were randomly assigned to one of four treatments: unmanipulated control (CN), oral secretion only (OS), mechanical damage only (MD), and mechanical damage plus oral secretion (MD+OS). A pilot study in 2022 (n = 6) did not contain the OS treatment but was included in the analysis. In total, 220 plants in 110 pots were used (treatment, n: CN, 29; OS, 23; MD, 29; MD+OS, 29).

To maximize emissions of HIPVs, we damaged the plants twice, once the evening before (17:00 h) and a second time the morning of the trial (7:00 h). The MD treatment consisted of using sand paper to abrade about 2 cm^2^ of the adaxial side of the two youngest developed leaves, avoiding the midvein. The OS treatment consisted of applying FAW regurgitant containing gut and salivary gland contents on the two youngest leaves (also on about 2 cm^2^) by gently squeezing the head of one caterpillar until it regurgitated. One caterpillar was used as a source of oral secretion for each pot. For the MD+OS treatment, the plants were mechanically damaged and immediately after, the oral secretion was applied directly on wounded tissue. After imposing the second damage treatment, approximately 14 hours after the initial treatment, one randomly selected live, 3^rd^ FAW or VAW caterpillar was pinned to the center of each leaf. Live caterpillars (as opposed to freeze-killed) were used to distinguish between scavenging and predatory behavior.

At 9:00 h, pots containing induced plants with pinned caterpillars were arranged in spatial blocks containing one treatment set (CN, OS, MS, MS+OS, where 1 treatment set = 1 n) in maize plots within the experimental field. Each pot was approximately 30 cm apart, forming a square with a random arrangement of treatments. Blocks were visited continuously over 75 minutes and the order in which sentinel caterpillars from the different treatments were removed in each block was recorded.

### Volatile Characterization

#### Plants and Insects

The same variety of maize plants used in the above field study obtained from a local famer were grown from seed at the University of Neuchâtel, Switzerland. Seeds were sown in individual plastic pots (4 cm x 10 cm) filled with commercial soil (Ricoter Erdaufbereitung, Aarberg, Switzerland). Potted plants were transferred to a glasshouse (L16:D8, T = 25°C ± 4, R.H.= 60-80%) and watered two to three times a week. After one week, fertilizer (Capito, Dotzigen, Switzerland) was added twice a week until use. Plants were used for experiments upon reaching two weeks of age.

Caterpillars of the fall armyworm *Spodoptera frugiperda* (J.E. Smith, 1797) (Lepidoptera; Noctuidae) were reared in a colony under quarantine conditions (T = 25°C ± 5, L16:D8, 60-80% R.H.) at the University of Neuchâtel (OFEV permit A140502). The colony was established from caterpillars collected at our field site during the 2023 field season and reared on the same artificial diet as the one used in the field experiment.

The same treatments imposed on the plants in the field were repeated for the volatile collection study with one exception, regurgitant from was collected from maize-fed 3^rd^ instar caterpillars in advance. Caterpillars were gently squeezed while on Parafilm (Sigma-Aldrich, Darmstadt, DE). The regurgitant was subsequently collected from the Parafilm, transferred to a microcentrifuge tube using a micropipette, and stored at −20°C until further use (about 1 week). For all treatments involving OS, 10 μl of regurgitant was applied using a micropipette.

#### Volatile Collection

We collected volatiles as described in Turlings et al (1998). Briefly, filtered air was pumped through a glass bottle volatile collection system containing the treated plants at a flowrate of 0.4L/min for two hours. Emitted volatiles were trapped on filters connected to this collection system (Porapak Q adsorbent, 25mg, 80-100, Merck, Darmstadt, DE). Each filter was then eluted with 150 μL of dichloromethane (Suprasolv, GC-grade; Merck, Darmstadt, DE) into glass vials to which 10 μL of internal standards ([20ng/μL] of n-octane and n-nonyl acetate) were added. Samples were then kept at −80°C until further analyses.

Volatile analyses were carried out using a gas chromatograph (Agilent 7890B) coupled to a mass spectrometer (Agilent 5977B). A 2 μL aliquot of each sample was injected in pulsed splitless mode onto a non-polar capillary column (Agilent HP-1; 30m length x 0.25mm ø x 0.25 μm thickness). After the injection, the temperature program maintained the temperature at 40°C for 3 minutes before increasing it progressively at a rate of 8°C min^−1^ until reaching 100°C; it thence increased it at 5°C min^−1^ until 200°C and subsequently rose it at 250°C for a 3 minutes post run. Volatiles were identified using the NIST mass spectral library and comparing retention times with those of commercial standards. Volatile emissions (ng hr^−1^) were calculated based on calibration curves of internal standards.

#### Statistical Analysis

All statical analyses were conducted in R (R version 4.2.1; R Core Team 2022). For the field study, we used binomial generalized linear models (GLMMs; glmmTMB, Brooks et al. 2017) to assess the effect of treatment on the first caterpillar attacked by predatory wasps. Date and plot ID were used as fixed and random variables, respectively. One-way analysis of variance (ANOVA) was used to assess the effect of treatment on the concentration of emitted volatile compounds. Volatile concentrations were log transformed to meet model assumptions. For all post hoc comparisons (emmeans; Length, 2021), the FDR correction method was used (Benjamini & Hochberg, 1995).

## RESULTS

### Sentinel Prey Experiment

Predatory social wasps quickly found, attacked, and removed sentinel caterpillars from maize plants. All replicate blocks were visited by wasps, resulting in at least one caterpillar attack and removal event per block. Nearly 45% of these attacks occurred within the first 30 minutes of the trials. The imposed treatments had a significant effect on the order in which caterpillars were attacked (Treatment: n = 3, *X^2^* = 27.738, P= <0.001). Caterpillars in the MD+OS, MD, and OS treatment were 11x, 5.4x, and 1.7x more likely to be attacked first compared to those in unmanipulated control plants, respectively (Fig. 1; Table S1). Of the observed attacks, all but one was perpetrated by *Polybia occidentalis* and *Polybia bribri* wasps. Both Polybia species were frequently found attacking the same caterpillar precluding the determination of which species arrived first. Though abundant in the field, only one caterpillar was removed by a *Polistes canadiensis* wasp.

**Fig 1.**
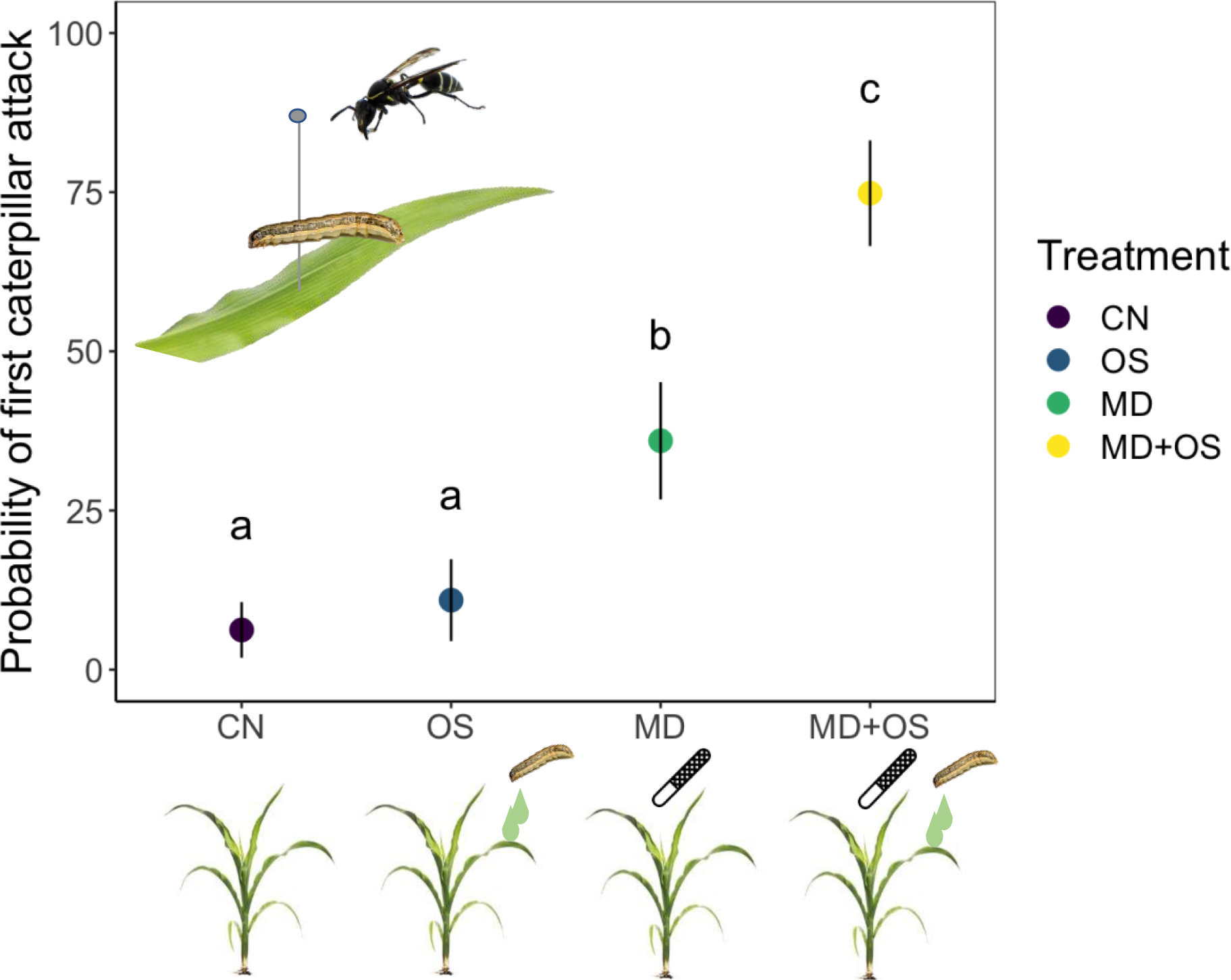
Marginal means and SE from a binomial GLMM from a sentinel prey removal experiment with maize plants subjected to one of four treatments. CN, OS, MD, and MD+OS represent the following treatments: control, FAW oral secretion (no damage), mechanical damage only, and the combination of mechanical damage and FAW oral secretion. The MD treatment was imposed using sandpaper, represented here with a file. Unique letters denote significant differences (P <0.05).

### Volatile Characterization

The imposed treatments quantitively and qualitatively affected the emission of maize seedlings (Fig. 2, Table S2, S3, and Fig. S3). Mechanically damaged (MD) plants and those which had oral secretion subsequently added to wounds (MD+OS), were associated with significantly increased emissions of several compounds, including (3E)-4,8-dimethyl-1,3,7-nonatriene (DMNT) and β-phenethyl compared to those treated only with oral secretions (OS) or unmanipulated controls (CN). Several other compounds (e.g., (E)-3-hexen-1-yl-acetate, β-linalool and indole) showed a similar pattern but were not significantly different after controlling for the family wise error rate (p > 0.05). The higher concentration of β-phenethyl and another homoterpene, (3E,7E)-4,8,12-trimethyl-1,3,7,11-tridecatetraene (TMTT), distinguished the MD+OS from the MD treatment. Interestingly, several unidentified compounds showed the opposite pattern. These constitutively emitted benzene derivatives were associated with a decreased emission upon damage.

**Fig 2.**
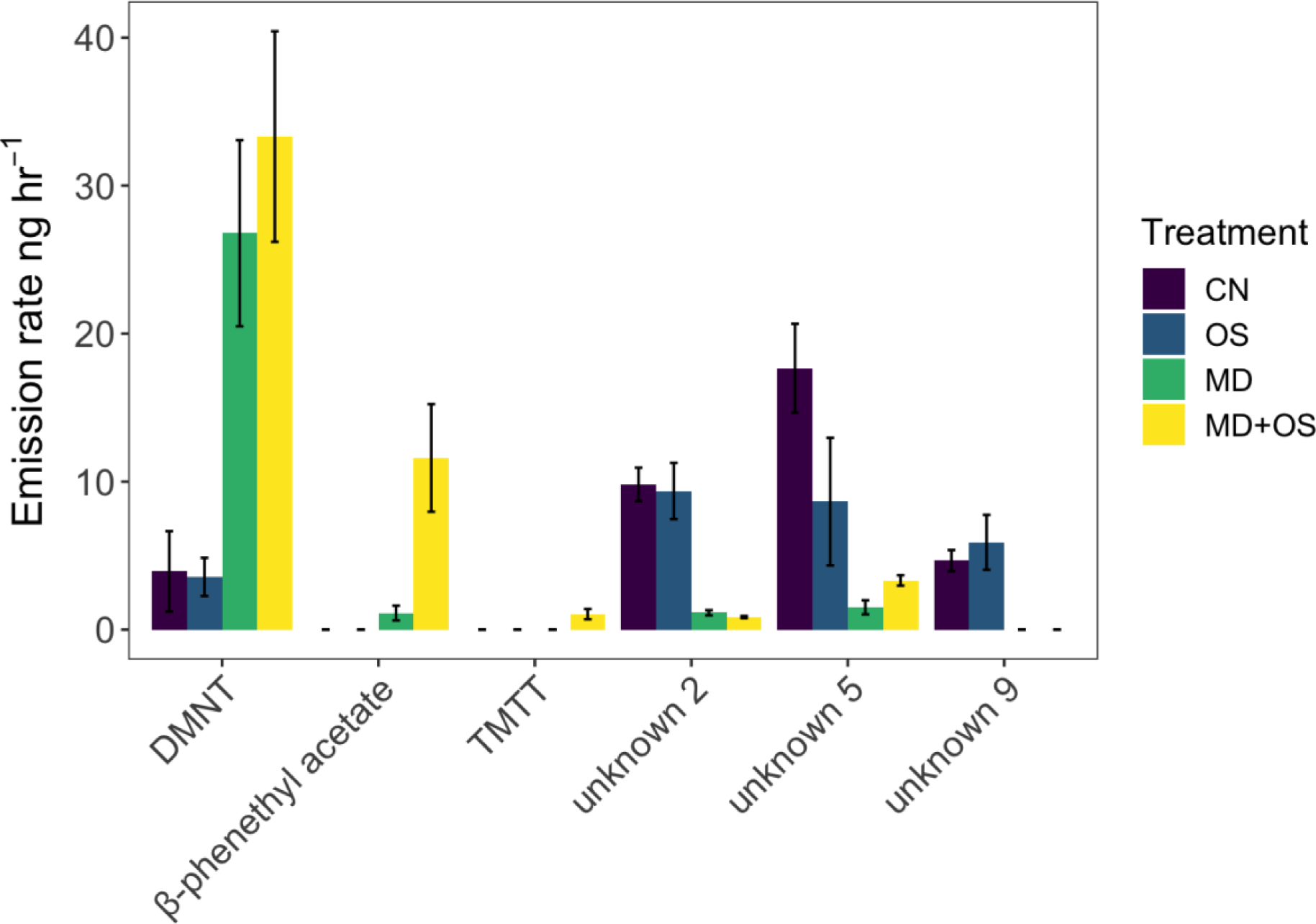
Concentration (ng hr^−1^) mean and ± SE of headspace volatile emissions from maize plants exposed to four treatments: an unaltered control (CN), FAW oral secretions (OS), mechanical damage (MD), and the combination of mechanical damage and FAW oral secretion (MD+OS).

## DISCUSSION

Since it was first proposed that herbivore-induced plant volatiles (HIPVs) contribute to the attraction of natural enemies (Dicke & Sabelis 1988; Turlings et al., 1990) numerous studies have confirmed this function, but comparatively few have been conducted under realistic field conditions (Dicke & Baldwin, 2010; M Heil, 2014; Kessler & Baldwin, 2003; Takabayashi & Shiojiri, 2019). While notable examples exist (De Moraes et al., 1998; Thaler, 1999; Poelman et al., 2009; Rodriguez-Saona et al., 2011; Schuman et al., 2012) the vast majority of studies have occurred under artificial conditions such as those typically imposed in flight tunnel and olfactometer assays. Such laboratory investigations are limited to assessing the attraction or repellence of predators and parasitoids towards an odor source, narrowing the inferences that can be drawn regarding predation and parasitism from resultant movement patterns in a more chemically complex environment (Roberts et al., 2023). Although informative, these experiments may overestimate the ecological relevance of HIPVs in tritrophic interactions as well as the regulatory capacity of natural enemies in pest control and the mitigation of yield loss (Poelman et al., 2009). This is especially true for parasitoids whose host will continue to feed despite being parasitized (Coleman et al., 1999). Field studies that attempted to address questions concerning the attraction of carnivorous arthropods and parasitic wasps to infested plants have commonly relied on natural herbivory for the production of attractive odors (Gomez & Zamora, 1994; Poelman et al., 2009). Although this approach is more realistic and enables the quantification of predation and parasitism, it fails to isolate factors contributing to the recruitment of natural enemies, making it challenging to evaluate the significance of key elements like the elicitors in the oral secretions of the attacking herbivore (Mamin et al., 2023; Waterman et al., 2019). In the present study, we demonstrated the relative importance of plant volatiles in prey location by predatory social wasp in an agricultural field, with the strongest attraction to and removal of sentinel caterpillars on maize plants with FAW oral secretion applied to mechanically inflicted wounds. This study marks the first instance of a study demonstrating the use of HIPVs by social predatory wasps in the field. Our findings not only represent a rare example validating the importance of HIPVs in prey location by these voracious predators under natural field conditions but also highlight the importance of insect-derived elicitors in mediating these interactions.

Compared to parasitic wasps, little is known about the role of plant volatiles in prey location for social wasps. Richter (1988) studied *Polybia* wasps, including *P. occidentalis* present at our field site, and found that foragers landed more frequently on leaves damaged by leafroller larvae than on undamaged leaves. Similarly, in a greenhouse experiment, Cornelius (1993) reported that naïve *Mischocyttarus flavitarsis* wasps captured more *Manduca sexta* and *Trichoplusia ni* caterpillars on tobacco plants with damaged leaves compared to those that were undamaged, strongly indicating a role of HIPVs. A more recent study using plants subjected to similar treatments as imposed here in a Y-tube olfactometer, found that *P. fastidiosucula* individuals preferred odors emitted from maize plants damaged either by FAW caterpillars or those with FAW oral secretions applied to mechanically damaged leaves (Saraiva 2017). The authors assessed the headspace volatiles and revealed that the two treatments produced volatile blends that were significantly different from control treatments and contained HIPVs known to attract parasitoids. In contrast to our findings, *P. fastidiosucula* showed no preference for mechanically damaged leaves compared to undamaged controls, indicating the importance of HIPVs rather than those that are immediately released upon tissue damage. The inconsistency between these studies suggests that specificity of the response needed to elicit searching behavior might be species-specific, as several other social wasps are known to be attracted to freshly cut leaves (Raw 1998). This inconsistency could also be explained by associative learning and differing levels of hunting experience of individuals between studies. Wasps, both parasitic (Giunti et al., 2015) and predatory (Elmquist & Landolt, 2018), are capable of associating cues with food rewards; HIPVs provide more reliable information regarding the likelihood of finding prey (Vet and Dicke, 1992). Over time, experienced individuals may become more discerning.

The characterization of headspace volatiles of maize plants strengthens the inference that the order in which caterpillars were attacked was a function of differing attraction to the odor blends resulting from the imposed treatments. Plants in the MD and MD+OS treatments showed increased emissions of several compounds, including β-phenethyl and two homoterpenes (DMNT and TMTT), compared to the OS and CN treatments. Consistent with other studies investigating HIPVs in maize, the MD+OS treatment showed the highest emissions of all three compounds shown to be inducible by *Spodoptera* caterpillars (D’Alessandro et al., 2009; Erb et al., 2015; Saraiva et al., 2017; Turlings & Tumlinson, 1992). Moreover, these induced compounds are known to attract parasitoids (D’Alessandro et al., 2009; Turlings et al., 1990) and one study indicated their attractiveness to one species of social wasp (Saraiva et al., 2017). The difference in emissions between the MD and MD+OS treatments was surprisingly minor in comparison with previous studies on inducible maize volatiles (Turlings et al., 1990; Turlings et al., 2002), but Mexican varieties are known to vary considerable in the amounts of HIPVs that they emit (Fritzsche-Hoballah et al, 2002). Overall, our findings suggest that the combination of wounding and oral secretions induces a specific volatile blend that either wasps have associated with food rewards or to which they are innately attracted, enhancing their ability to locate and capture prey.

While we predicted differences between the volatile blends associated with the four treatments, we did not expect to find a decrease in emission of several constitutively emitted compounds upon damage (Fig S1) these compounds have not previously been reported for maize and remain to be identified. Indeed, the downregulation of volatile compounds is rarely reported. A study by (Peñaflor et al., 2011) found that oviposition by FAW on maize plants was associated with a decreased emission of constitutively emitted volatiles as well as a suppression of those that are induced upon herbivory. Suppression of plant defenses by insect eggs has also been reported for *Arabidopsis* (Bruessow et al. 2010), although the opposite may also be the case (Stahl et al., 2023). In our case, we observed decreased emission of constitutive compounds in the MD as well as the MD + OS treatment. Consequently, insect-associated suppression factors do not explain this result. Further work with this particular and related maize variety is necessary to understand the causes and consequences of this unusual finding.

Besides confirming, to our knowledge for the first time in nature, the importance of elicitor-trigger volatile emissions for the attraction of the third tropic level, this study also is of applied relevance. The hunting proficiency of social wasps has been well-documented, particularly in their pursuit of lepidopteran pests, prompting initial field trials exploring their potential as biocontrol agents (Institute of Agricultural and Forestry Sciences of Shang-Chiu, 1976; Rabb & Lawson, 1957). With an increased awareness of the adverse effects of agrochemicals on human health and the environment, there is a growing need for sustainable pest management strategies (Barzman et al., 2015). This is particularly true for the control of FAW which, upon its recent invasion of Africa and Asia, has led to an unprecedented increase in the use of insecticides (Kenis et al., 2022; Yang et al., 2021).

Consequently, there is renewed interest in considering social wasps as potential biocontrol agents, especially against FAW (Montefusco et al., 2017; Prezoto et al., 2019a; Saraiva et al., 2017; Southon et al., 2019). In addition to reinforcing the need for this field of research, our study emphasizes the significance of plant volatiles likely induced by elicitors in oral secretions of FAW for attracting these effective predators. This new understanding can provide the basis for the development of novel crop protection methods, such as utilizing chemical elicitors to enhance crop plants’ attractiveness to biocontrol agents, including social wasps (Sobhy et al., 2014; Turlings & Erb, 2018).

## ACKNOWLEDGMENTS

We would like to thank Alfredo Lopez-Rojas and Raul González-Salas for their assistance with fieldwork and Carla Cristina Marques Arce for her expertise with the collection and analysis of plant volatiles.

