## supplemental material for "Field evidence for the role of plant volatiles induced by caterpillar-derived elicitors in the prey location behavior of predatory social wasps"

Table S1. Post hoc pairwise comparisons resulting from the GLMM of the sentinel prey removal study. P-values calculated from estimated marginal means were adjusted for multiple comparisons using the FDR method.

| Contrast | Estimate | SE | Z ratio | P value |
| --- | --- | --- | --- | --- |
| CN - OS | -0.611 | 0.966 | -0.633 | 0.527 |
| CN - MD | -2.133 | 0.831 | -2.568 | 0.015 |
| CN - (MD+OS) | -3.801 | 0.86 | -4.418 | 0.000 |
| OS - MD | -1.522 | 0.742 | -2.053 | 0.048 |
| OS - (MD+OS) | -3.19 | 0.78 | -4.089 | 0.000 |
| MD - (MD+OS) | -1.668 | 0.586 | -2.846 | 0.009 |

Table S2. Retention time (RT) and mean ± SE of volatile compounds (ng hr^-1^) emitted from maize plants subjected to four treatments:

control (CN; n=5), oral secretion (OS; n=6), mechanical damage (MD, n=5), and mechanical damage with oral secretions (MD+OS).

| **RT** | **Compound** | **CN** | | | **OS** | | | **MD** | | | **MD+OS** | | |
| --- | --- | --- | --- | --- | --- | --- | --- | --- | --- | --- | --- | --- | --- |
| 10.28 | (E)-3-Hexen-1-ol-acetate | 0.36 | ± | 0.16 | 0.39 | ± | 0.21 | 2.74 | ± | 0.76 | 3.90 | ± | 2.09 |
| 12.31 | β-Linalool | 2.59 | ± | 1.14 | 3.78 | ± | 1.11 | 13.30 | ± | 5.57 | 14.40 | ± | 4.12 |
| 12.69 | DMNT^1^ | 3.94 | ± | 2.71 | 3.56 | ± | 1.29 | 26.78 | ± | 6.29 | 33.31 | ± | 7.11 |
| 15.16 | 1,8-nonadien-3-ol | 0.00 | ± | 0.00 | 0.00 | ± | 0.00 | 0.23 | ± | 0.18 | 0.91 | ± | 0.39 |
| 16.03 | β-Phenethyl acetate | 0.00 | ± | 0.00 | 0.00 | ± | 0.00 | 1.12 | ± | 0.50 | 11.60 | ± | 3.64 |
| 16.94 | Indole | 0.99 | ± | 0.49 | 1.87 | ± | 1.01 | 6.33 | ± | 1.85 | 6.14 | ± | 1.84 |
| 17.62 | unknown 1 | 5.02 | ± | 0.63 | 5.14 | ± | 0.94 | 0.98 | ± | 0.26 | 0.97 | ± | 0.13 |
| 17.71 | unknown 2 | 9.81 | ± | 1.13 | 9.36 | ± | 1.90 | 1.14 | ± | 0.17 | 0.85 | ± | 0.07 |
| 17.90 | unknown 3 | 10.37 | ± | 1.14 | 9.91 | ± | 2.09 | 0.26 | ± | 0.16 | 0.24 | ± | 0.08 |
| 18.38 | unknown 4 | 11.24 | ± | 2.75 | 5.49 | ± | 3.34 | 0.33 | ± | 0.20 | 1.41 | ± | 0.16 |
| 18.88 | unkown 5 | 17.66 | ± | 3.00 | 8.65 | ± | 4.31 | 1.51 | ± | 0.48 | 3.32 | ± | 0.35 |
| 19.67 | uknown 6 | 2.46 | ± | 0.44 | 2.44 | ± | 0.73 | 0.18 | ± | 0.18 | 0.00 | ± | 0.00 |
| 19.83 | uknown 7 | 6.33 | ± | 1.94 | 1.75 | ± | 1.05 | 0.51 | ± | 0.17 | 0.73 | ± | 0.28 |
| 20.41 | β-Bergamotene | 1.27 | ± | 0.39 | 2.32 | ± | 1.53 | 1.93 | ± | 0.57 | 2.29 | ± | 1.26 |
| 20.86 | (E)-β-Farnesene | 0.64 | ± | 0.32 | 1.81 | ± | 1.43 | 1.68 | ± | 0.70 | 2.46 | ± | 1.29 |
| 22.16 | uknown 8 | 0.09 | ± | 0.09 | 1.44 | ± | 1.01 | 3.40 | ± | 1.05 | 2.26 | ± | 0.36 |
| 23.03 | uknown 9 | 4.66 | ± | 0.71 | 5.90 | ± | 1.85 | 0.00 | ± | 0.00 | 0.00 | ± | 0.00 |
| 23.16 | uknown 10 | 2.17 | ± | 0.38 | 2.93 | ± | 0.91 | 0.00 | ± | 0.00 | 0.00 | ± | 0.00 |
| 23.74 | TMTT^2^ | 0.00 | ± | 0.00 | 0.00 | ± | 0.00 | 0.00 | ± | 0.00 | 1.04 | ± | 0.34 |
|  | Total | 96.66 | ± | 9.72 | 81.48 | ± | 9.39 | 74.45 | ± | 15.05 | 100.12 | ± | 17.15 |

^1^ (E)-3,8-dimethyl-1,4,7-nonatriene (DMNT)

^2^ (E,E)-4,8,12-trimethyltrideca-1,3,7,11-tetraene (TMTT)

Table S3. P-values from One-Way ANOVA to assess the overall differences among four treatments imposed on maize plants and those from post hoc pairwise comparisons performed using estimated marginal means. P-values were adjusted for multiple comparisons using the FDR method for both the overall ANOVA (n=number of compounds assessed) and pairwise comparisons. CN, OS, MD, and MD+OS treatments represent the following treatments: control (n=5), oral secretion (n=6), mechanical damage (n=5), and mechanical damage with oral secretions (n= 6). Emboldened text represents significant differences (p <0.05).

| **Compound** | **One Way ANOVA** | **CN - MD** | **CN - (MD+OS)** | **CN - OS** | **MD - (MD+OS)** | **MD - OS** | **(MD+OS) - OS** |
| --- | --- | --- | --- | --- | --- | --- | --- |
| E-3-Hexen-1-ol-acetate | 0.389 | - | - | - | - | - | - |
| β-Linalool | 0.124 | - | - | - | - | - | - |
| DMNT^1^ | **0.002** | **0.008** | **0.003** | 0.981 | 0.981 | **0.009** | **0.003** |
| 1,8-nonadien-3-ol | 0.420 | - | - | - | - | - | - |
| β-Phenethyl acetate | **0.000** | 0.480 | **0.000** | 1.000 | **0.005** | 0.480 | **0.000** |
| Indole | 0.968 | - | - | - | - | - | - |
| unknown 1 | **0.000** | **0.000** | **0.000** | 1.000 | 1.000 | **0.000** | **0.000** |
| unknown 2 | **0.000** | **0.000** | **0.000** | 0.931 | 0.931 | **0.000** | **0.000** |
| unknown 3 | **0.000** | **0.000** | **0.000** | 1.000 | 1.000 | **0.000** | **0.000** |
| unknown 4 | **0.002** | **0.000** | **0.006** | 0.099 | 0.314 | **0.024** | 0.314 |
| unkown 5 | **0.001** | **0.000** | **0.003** | **0.055** | 0.261 | **0.018** | 0.343 |
| uknown 6 | **0.002** | **0.006** | **0.003** | 0.966 | 0.966 | **0.008** | **0.003** |
| uknown 7 | 0.080 | - | - | - | - | - | - |
| β-Bergamotene | 1.000 | - | - | - | - | - | - |
| (E)-β-Farnesene | 1.000 | - | - | - | - | - | - |
| uknown 8 | 0.137 | - | - | - | - | - | - |
| uknown 9 | **0.000** | **0.000** | **0.000** | 1.000 | 1.000 | **0.000** | **0.000** |
| uknown 10 | **0.001** | **0.002** | **0.002** | 1.000 | 1.000 | **0.002** | **0.002** |
| TMTT^1^ | **0.008** | 1.000 | **0.004** | 1.000 | **0.004** | 1.000 | **0.004** |

^1^ (E)-3,8-dimethyl-1,4,7-nonatriene (DMNT)

^2^ (E,E)-4,8,12-trimethyltrideca-1,3,7,11-tetraene (TMTT)

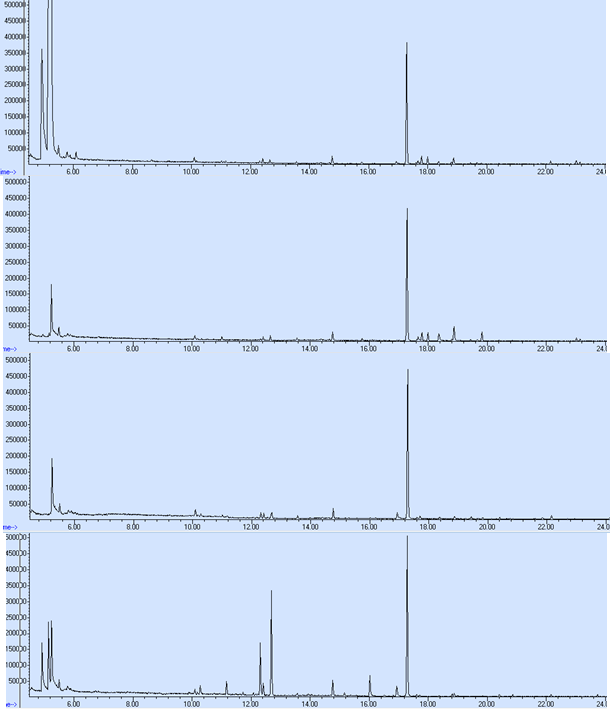

CN

OS

MD

MD + OS

a

b

c

d

e

f

g

h-j

k

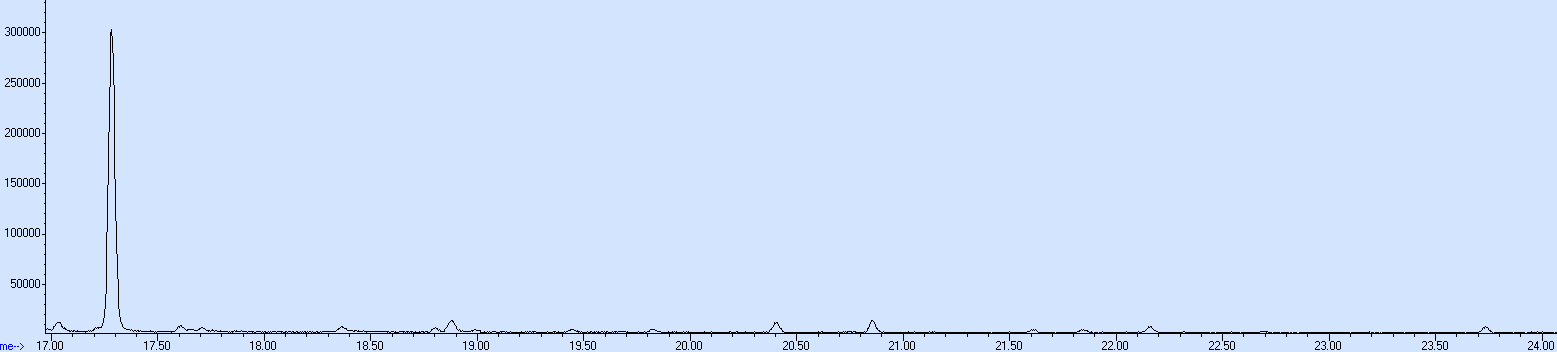

g

p

l

m

n

C

B

A

^1^ (E)-3,8-dimethyl-1,4,7-nonatriene (DMNT)

| Letter | RT | Compound |
| --- | --- | --- |
| a | 10.28 | (E)-3-Hexen-1-ol-acetate |
| b | 12.31 | β-Linalool |
| c | 12.69 | DMNT |
| d | 15.16 | 1,8-nonadien-3-ol |
| e | 16.03 | β-Phenethyl acetate |
| f | 16.94 | Indole |
| g | 17.27 | Internal Standard |
| H | 17.62 | Unknown 1 |
| i | 17.71 | Unknown 2 |
| J | 17.91 | Unknown 3 |
| k | 19.83 | Unknown 4 |
| l | 20.41 | β-Bergamotene |
| m | 20.86 | (E)-β-Farnesene |
| n | 22.16 | Unknown 8 |
| p | 23.74 | TMTT |

^2^ (E,E)-4,8,12-trimethyltrideca-1,3,7,11-tetraene (TMTT)

Fig S2. Representative chromatograms (A,B) of identified and unidentified compounds and retention times (C) for maize plants subjected to four treatments, unmanipulated control (CN), FAW oral secretion (OS), mechanical damage (MD), and the combination of OS and MD (MD + OS). A few peaks with later retention times were minor including TMTT, which was only present in the MD + OS treatment (B). Note that the unknown compounds listed appear to be constitutively emitted with the cessation of emission upon wounding.
